# From video to behaviour: an LSTM-based approach for automated nest behaviour recognition in the wild

**DOI:** 10.1101/2024.10.25.620052

**Authors:** Liliana R. Silva, André C. Ferreira, Irene Martínez-Baquero, Arlette Fauteux, Claire Doutrelant, Rita Covas

## Abstract

Studies of animal behaviour usually rely on direct observations or manual annotations of video recordings. However, such methods can be very time-consuming and error-prone, leading to sub-optimal sample sizes. Recent advances in deep-learning show great potential to overcome such limitations, nevertheless, most currently available behavioural recognition solutions remain focused on captivity settings.
Here, we present a deployment-focused framework to guide researchers in building behavioural recognition systems from video data, using Long Short-Term Memory (LSTM) networks to classify behavioural sequences across consecutive frames.
LSTMs allowed to: 1) monitor nest activity by detecting the birds’ presence and simultaneously classifying the type of trajectory: i.e., nest-chamber entrance or exit; and 2) identify the behaviour performed: building, aggression or sanitation. Using our framework, we outperformed human annotators when jointly considering error and speed. Model performance improved with challenging training instances, and remained robust even with modest sample sizes. LSTM also outperformed YOLO (“You Only Look Once”), highlighting the critical role of temporal sequence information in behavioural analysis.
We demonstrate that our approach is replicable across three bird species and applicable to deployment videos, highlighting its value as a generalizable and transferable tool for long-term studies in the wild.

**DATA AVAILABILITY:** Scripts, models, and data required to reproduce this work are available on Zenodo (DOIs: 10.5281/zenodo.18681623 and 10.5281/zenodo.18695178).

## INTRODUCTION

Animal behaviour studies frequently rely on video for behavioural characterisation and quantification (Anderson & Perona, 2014). However, behaviour analysis through video is costly given the complexity and variable nature of behavioural data and the ever-growing size of longitudinal datasets. Video analysis requires expertise, annotators, or commercial software, demanding significant resources, whereas manual coding is subjective, error-prone and tedious (Anderson & Perona, 2014). Despite these drawbacks, video still remains one of the most effective tools for behavioural data-collection.

There is increasing effort to automate behavioural annotation by applying machine-learning algorithms to human and animal behavioural datasets, with recent emphasis on the use of deep-learning techniques (Christin et al., 2019). These models allow for automatic detection and extraction of features, often surpassing human capabilities (Pichler & Hartig, 2023). Most recent advances on behavioural automation from video, either open-source (e.g., Harris et al., 2023) or commercial hardware/software (e.g., Ethovision), are focused on captivity settings (reviewed in Panadeiro et al., 2021; but see e.g., Mounir et al., 2023). Although they constitute important advances, such frameworks lack applicability for contexts characterised by varying recording conditions, mutable environment and unrestricted movement, such as seen in the wild.

Automated video-based behavioural analyses can be performed using a single frame (e.g., Norouzzadeh et al., 2018) or a sequence of frames (e.g., Williams & DeLeon, 2020). Single-frame approaches risk missing the dynamic nature of behaviour, while multi-frame methods better capture changes in position and pose over time (e.g., Bohnslav et al., 2021). For example, a bird flying toward a nest may appear to be entering in one frame, but only a sequence reveals whether it actually does. Behavioural frame sequences can be analysed using either body coordinates mapped to behavioural descriptors (e.g., Mathis et al., 2018) or automatically extracted features (e.g., Harris et al., 2023). In both cases, these representations can be modelled with temporal networks to capture sequential dependencies for behavioural classification (e.g., Williams & DeLeon, 2020), simultaneously detecting the animals and respective behaviour. While such approaches have proven to be successful for behavioural analyses, they have been mostly applied to humans (e.g., Van Houdt et al., 2020), constrained settings (e.g., Hu et al. 2023) or data types other than video (e.g. accelerometery: Mao et al., 2023, but see Williams & DeLeon, 2020) leaving a gap of temporal modelling application for behaviour in wild from video data.

Deep learning is now widely accessible, enabling numerous proof-of-concept works but arguably fostering over-optimism about its application (Saidi et al., 2024). The deployment of deep-learning models is challenging, especially for a realistic application in the wild and long-term as models that perform well during development might perform poorly when taken to the real-world (see Wilson et al., 2025). This can be a result of multiple factors from poorly designed training and testing datasets to unexpected events in the wild or insufficient understanding of deployment contexts. Furthermore, despite the abundance of modelling options, there remains a gap in their practical real-world application and in guidance on how to appropriately extract, handle and train (behavioural) data that will align with temporal modelling and ecological questions.

Here, we developed a temporal automation framework, based on Long Short-Term Memory networks (“LSTM”, Hochreiter & Schmidhuber, 1997), with the aim of guiding researchers in building their own behavioural recognition system directly from video data, using open-source Python libraries (Fig. 1A), with a focus on realistic, long-term applications in the wild. Our framework builds on a long-term study of the sociable weaver (*Philetairus socius*), based on complex behavioural data manually collected since 2014, and offers a basis for models that can be extended to other systems.

**Figure 1.**
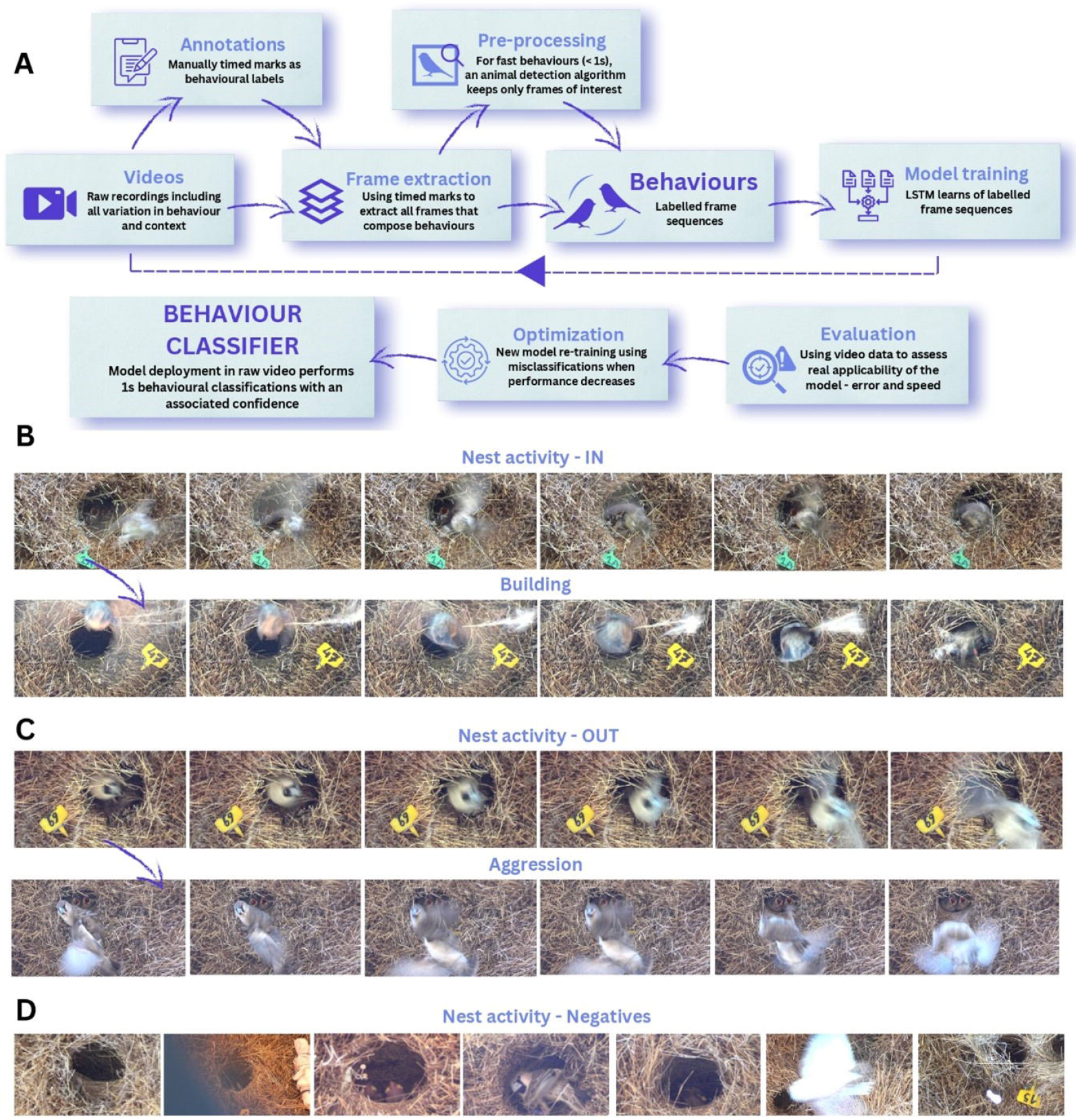
(A) Pipeline for automatic behavioural classification from video using manually annotated data with LSTMs. Frame sequences showing hierarchical behaviour detection in sociable weavers: (B) “IN” (entrance) with building when entering with straw; (C) “OUT” (exit) with aggression when expelling another bird. (D) Single-frame negative class examples used to improve model efficiency.

We first describe model development, including considerations related to the selection of challenging training instances, highly unbalanced behavioural samples, variations in behaviour duration, and the impact of dataset size on model learning. We then evaluate our approach by: 1) assessing the performance and speed of the model for behaviour classification, 2) comparing model performance to human analysers and 3) assessing the model’s ability to output data that could detect real biological effects. In addition, we compare our multi-frame approach, which captures the temporal dynamics of behaviour, with a single-frame model, YOLO (“You Only Look Once”, Redmon et al., 2015), a convolutional network that classifies images independently, and which has been recently used for video-based behaviour recognition (e.g., Chan et al., 2025), appealing for its simplicity. To assess the generalisation capabilities approach, we use our framework to analyse the nesting behaviour of two other species of birds: blue (*Cyanistes caeruleus*) and great tits (*Parus major*). Finally, we discuss some of the caveats of moving from proof-of-concept to model deployment.

## METHODS

### Behavioural data

Between 2014 and 2021, videos of sociable weavers were collected by field assistants in the wild in South Africa. We standardised recordings, with each nest chamber filmed with fixed cameras in Full HD quality (n= 1239 nests, 5,143 2h-videos, 208,548 bird visits). We extracted temporal-preserved frames from the manually annotated raw videos and constructed separate datasets for each type of behaviour (i.e., nest activity, building and aggression). We manually annotated behaviours per second, through a pre-processing step to keep the specific frames containing the behaviour. All details for data collection, manual annotation, quality control, frames’ extraction and pre-processing are in the Supplementary Material, hereafter referred to as “SM”, S1:S6, and illustrated on Fig.1A).

### Behavioural classification

For behaviour classification of sociable weavers, we identified four distinct behaviours (Fig. 1B–1C): entering the nest chamber (“entrance”), used as a proxy for nest provisioning; exiting the nest chamber (“exit”); bringing straws to the nest chamber for construction (“building”); and exhibiting aggression by expelling a conspecific from the nest chamber (“aggression”). These four behaviours only make a minuscule part of all recording time (<1%). The remaining time is hereafter referred to as “negative class” (“NC”, Fig. 1D). We targeted NC frame sequences that are challenging to classify, (hereafter hard negatives) because they are likely to be confused with the focal behaviours (e.g., distinguishing nest fly-bys from entrances); Fig. 1D, excluding the leftmost example), rather than randomly sampling non-behavioural segments, which would yield mostly easy classifications (e.g., static nest chambers). To quantify their impact, we compared model performance with varying proportions of hard negatives (10% vs. 55% of selected hard negatives; see SM-S6E for modelling details and results).

Additional challenges arise from strong class imbalance, as entrances and exits comprise the vast majority of behavioural frames, whereas building (6.5%) and aggression (0.16%) are rare. Building and aggression are also context-dependent: building occurs only when birds enter with straw in their bills, while aggression is consistently followed by the aggressed bird leaving. In addition, aggression events are longer (≈16 frames vs. ≈6 for other behaviours), reflecting the fact that human observers typically require more frames to recognise them.

To address these differences, we implemented a hierarchical framework consisting of three models: (i) nest activity (entrance vs. exit vs. NC), (ii) building (entrance with straw vs. entrance without straw), and (iii) aggression (exit with aggression vs. exit without aggression). The first model, trained on a larger dataset, screens the entire video for nest activity, whereas the latter two, trained on smaller datasets, make predictions only within events identified as entrance (building) or exit (aggression). To further accommodate differences in behaviour duration, all three models used six-frame input sequences: frames were consecutive for the nest activity and building models, while the aggression model used six frames sampled every third frame, spanning 16 frames in total (i.e., frames 1, 4, 7, 10, 13, and 16).

All models combined a pre-trained VGG19 backbone (without the final classification layer) for frame-level feature extraction with an LSTM layer to capture temporal dependencies. Features were further reduced by four dense layers, and classification was performed using a softmax layer for nest activity (entrance, exit, NC) or a sigmoid layer for building (entrance with straw vs. entrance without) and aggression (OUT with aggression vs. OUT without). Transfer learning was applied by initialising the nest activity model with ImageNet weights (following a previous classification task in this species; Ferreira et al., 2020), and subsequently using the trained nest activity model to initialise the building and aggression models, given their closer similarity. All training details can be found on SM-S5.

Because model performance can depend on training dataset size, we evaluated the error rates of models developed for nest activity detection using different dataset size (25%, 50%, 75%, and 100% of the data; see SM-S6F for modelling details and results).

Finally, we compared our framework with a single-frame approach based on YOLOv8 (Ultralytics, 2023) for nest activity detection using equivalent datasets (see SM-S6G for modelling details and results).

### Evaluation

We first evaluated model accuracy using balanced validation datasets (i.e., equal class sizes, Table 1). To test real-world applicability and generalisation, models were evaluated on manually annotated full video recordings, matching those used in future deployment (“deployment videos”), and reserved solely for final testing. Performance was measured in terms of error and processing speed relative to humans, using double-screened annotations and average human error serving as reference benchmarks (see SM-6 for details on this evaluation, Table 1). Error was quantified as false positives (FP; behaviours annotated without relevant activity) and false negatives (FN; missed behaviours of interest), over the number of actual visits per video. These measures directly capture deployment implications: a 100% FP rate means all predictions are wrong, while a 100% FN rate means all real behaviours are missed. Complementary evaluation metrics are provided in the SM-6D. In addition, nest activity and building models were biologically validated by testing models’ predictions that could be used to directly test hypotheses with *a priori* known outcomes: 1) that age and number of nestlings are positively related to nest activity and 2) building activity is related to the nest stage (higher during incubation). For aggression, no a priori predictions were available. All details of models’ evaluation are provided in SM-S6.

**Table 1.**
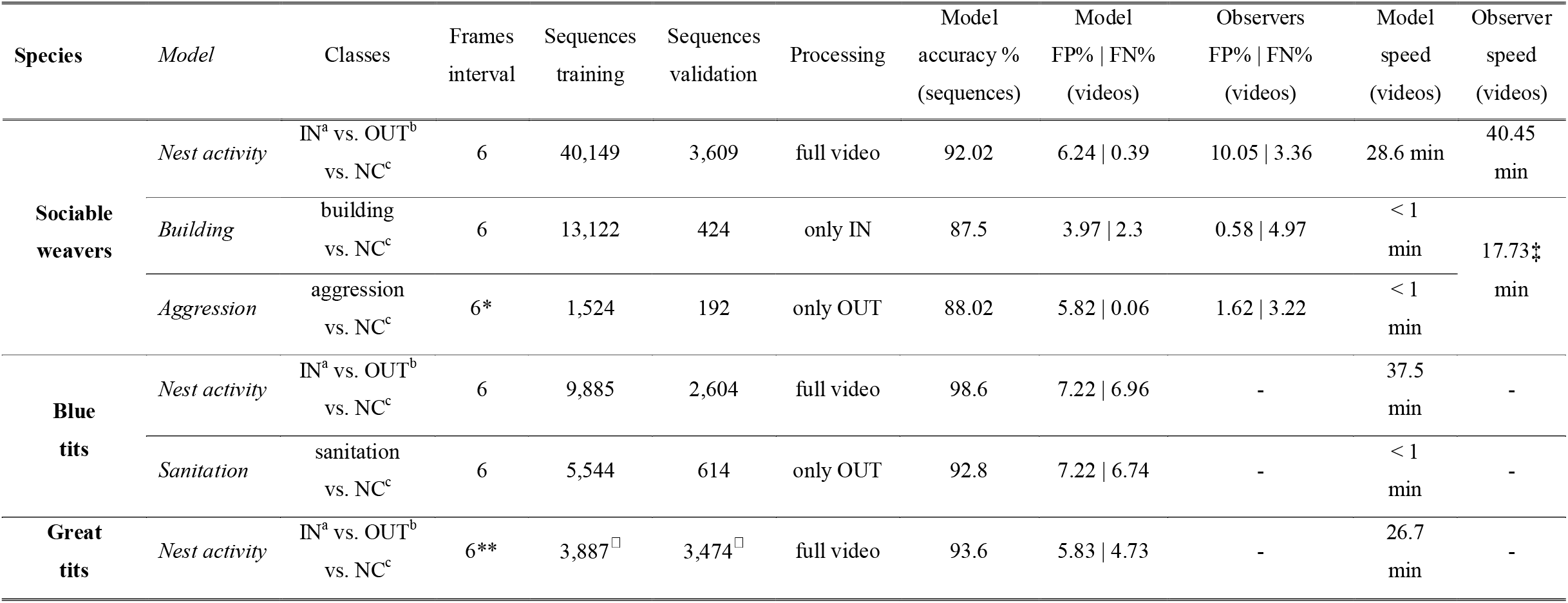
Behavioural detection models’ training details and performance. Training details include: classes, interval of frames, training and validation datasets Performance was assessed as i) the model frame-accuracy obtained at the validation stage, but also, ii) using videos (2h) to assess the produced false ves (“FP”) and false negatives (“FN”), as well as speed (i.e., annotation time), performed by model and observers, to assess deployment success. ^a^Next-chamber entrance, ^b^nest-chamber exit, ^c^negative class, *every three frames, **every six frames, ^□^“leave-one-out” approach (SM-8D), ‡performed simultaneousy

### Species and context generalisation

To further test the generalisability of our framework, we automated behaviour detection of two additional species, monitored in 2024, as part of two long-term studies: blue tits in Corsica (France) and great tits in Wytham Woods (UK). In both cases, the breeding pairs were recorded in settings that differed remarkably from that of sociable weavers. These recordings were made from inside the nest box using an embedded camera positioned to record the entrance, depicting the provisioning of nestlings. Blue tit recordings were collected continuously (n=17 nest-boxes, 185 24h-videos), capturing the activity inside the nest, including sanitation events (i.e., when a bird carries a faecal bag outside). By contrast, great tit recordings (n=15 nest-boxes, 38 2h-videos) were filmed from the back of the nest-box and limited mostly to distinguishing arrivals and departures. Building on the approach developed for sociable weavers, we automated the detection of entrance and exit for both tit species, and sanitation for blue tits only (Table 1, Fig. 2, SM-S7:S8 for additional details).

**Figure 2.**
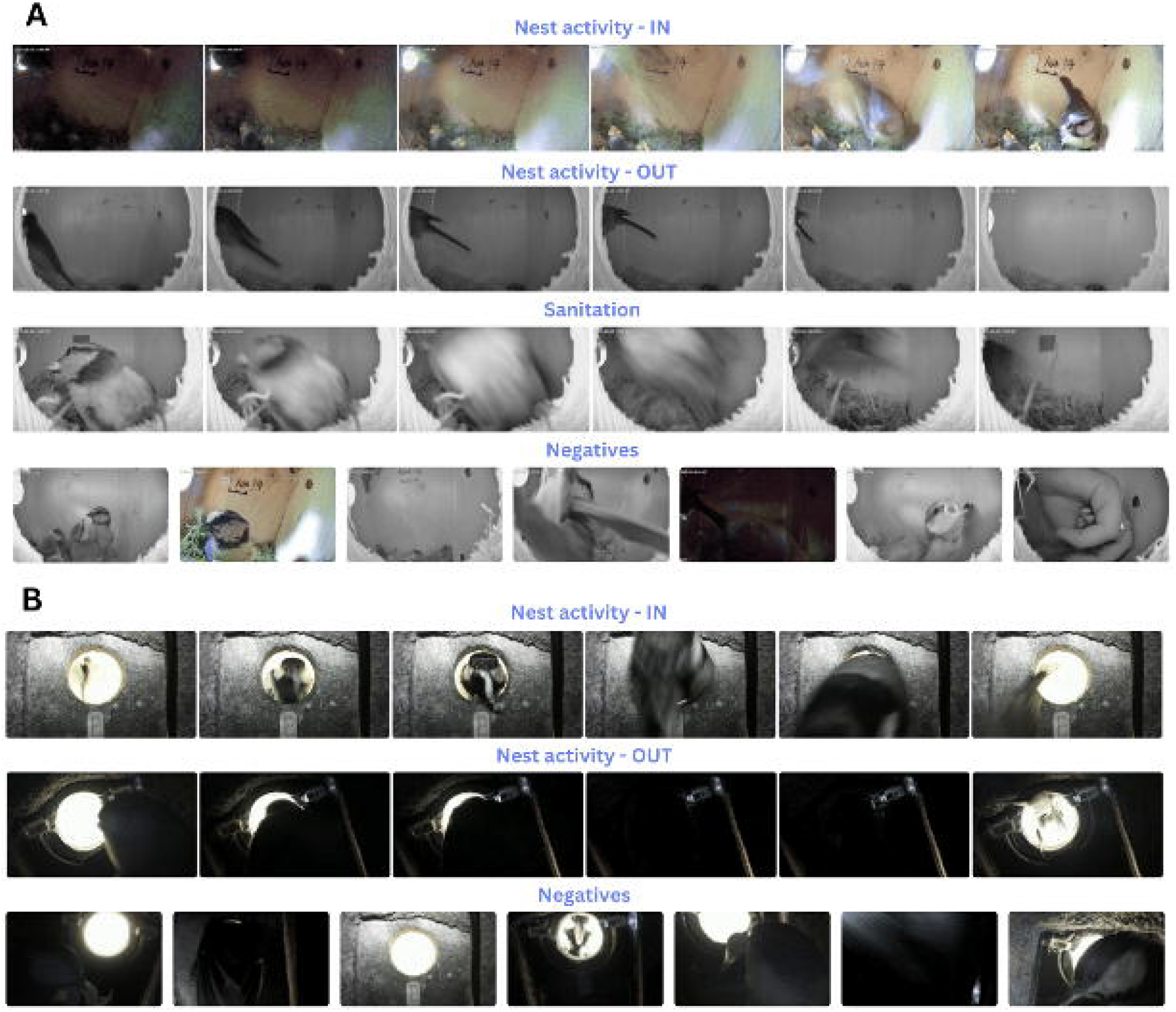
Frame examples of automated nest-box activity detection in (A) blue tits and (B) great tits, with corresponding negative classes. In blue tits, sanitation was hierarchically detected on exit.

## RESULTS

In the sociable weaver, all models had a validation accuracy >87.5% (Table 1). FP and FN were overall lower than those of annotators, while all models outperformed them in analysis speed (Table 1). Automated data allowed for the detection of biologically significant effects previously known for sociable weavers (details and results in SM-S6A:B, Fig.S2-S3).

Using nest activity automation the sociable weavers, we found that: (1) including hard negatives during training had a strong effect, reducing error by ∼60% (from 19% FP and 1.02% FN to 7.67% FP and 0.96% FN, SM-S6E); (2) overall training dataset size was not limiting, as even 25% of the original dataset (10,037 sequences) yielded error rates close to those obtained with the full dataset (SM-S6F); and (3) adopting a single-frame YOLOv8 approach markedly worsened performance, increasing error nearly fourfold (from 7.4% FP and 6.84% FN with LSTM on a comparable dataset to 54.7% FP and 1.06% FN using YOLOv8, SM-S6G).

Our framework was successfully applied to the behaviours of two tit species, also achieving high accuracy, speed, and low detection error (Table 1, modelling details in SM-S7:S8).

## DISCUSSION

With this work, we present the deployment of a powerful behavioural analysis framework that automates the identification of nest behaviours. We show that temporal modelling, through LSTMs, is effective for automating behaviour classification and enables researchers to leverage their own previously annotated data, often already existent in long-term projects. While the behaviours automated in this work were challenging to classify, our models matched experienced annotators (with over six months of practice) and outperformed inexperienced ones (see SM-S6A:B for error rates by experience). Additionally, our framework, coupled with the 24/7 working time of a computer, allowed us to increase by eight times the processing speed analyses (from analysing 41.25 to 345.68 videos per week, see SM-S6C). The effectiveness of this framework is further demonstrated by its current application in the sociable weaver long-term project, where it reduced manual effort by over 2,600 working hours across four recording field seasons. Additionally, we demonstrate that the approach developed here can be readily applied to other wild birds, being implemented in other long-term studies.

Dataset quality and composition are key factors when training deep-learning models (Gong et al., 2023). Reducing the sociable weavers’ nest activity dataset to ∼10,000 behavioural sequences (25% of the original size; SM-S6F) had little effect on LSTM performance. However, when most of the negative instances were easy classification examples, the error doubled on deployment videos, even though model train-validation metrics were similar for models with low and high hard negatives (accuracies: 97.30% and 94.44%, respectively, SM-S6E). These results highlight the need for high-quality training datasets and evaluation on deployment videos to ensure robust real-world performance. Therefore, here we provide the largest annotated dataset for nest-related behaviour to date, compiled over several years by multiple annotators and capturing wide natural variability, which is essential for robust generalisation (SM-S9). By contributing to the growing body of behavioural annotation datasets (e.g., MammalNet, Chen et al., 2023) and providing a rare large-scale dataset on bird behaviour, our work facilitates cross-species comparisons, benchmarking, and transfer learning.

Comparing YOLO, a single-frame approach previously shown to perform well for behavioural classification (Chan et al., 2025), with temporal modelling, we found that YOLO performed substantially worse. This was expected, since behaviours are defined by actions unfolding over time (Bohnslav et al., 2021). Although both models showed promising train–validation metrics (accuracies: 96.11% for LSTM, 80.56% for YOLO; SM-S6G), the number of produced false positives during deployment differed drastically: from 960 real visits, YOLO predicted 5,990, whereas the LSTM model predicted 932. The LSTM slightly underestimates visits due to missed detections but remains close to the real scenario, whereas YOLO drastically overestimates by marking continuously and producing correct detections amid numerous false positives. Nonetheless, single-frame methods may still succeed when a specific pose/appearance uniquely indicates a behaviour. Additionally, although our specific LSTM-based model proved to be highly effective in our case, we note that the rapidly expanding field of machine learning yields powerful alternative approaches (e.g. Marks et al., 2022; Sun et al., 2023). With growing number of large behavioural datasets like ours, future work should further investigate which approach is more suitable across contexts.

Although useful, automation of behavioural analyses through deep-learning, especially for video, has limitations. First, model performance can decline over time affecting deployment. Specifically, factors associated with long-term studies, such as systematic changes in recording conditions and study design, can introduce unexpected variation not included in the training, reducing model performance. Therefore, for long-term deployments, deep-learning models require ongoing monitoring and optimisation (Fig.1A). Monitoring models’ performance involves regular inspection of models’ outputs and reusing model mispredictions to re-train models with new “harder” examples. This process enhances the models’ generalisation and longevity—key objectives for researchers using deep-learning.a In addition, selecting, developing and deploying such models takes time (ranging from 3-6 months depending on the complexity), highlighting the importance of creating generalisable and transferable automation approaches that can be shared across research projects, as was done here. Finally, this automation work focused on short (<1s), dynamic, non-state complex behaviours, but further research should test whether LSTMs can extend to longer behaviours.

## ACKNOWLEDGEMENTS

We thank several researchers, field team managers, assistants and volunteers for help with data collection, management and analysis from the Sociable Weaver, the Corsican UQUAM-CEFE Tit and Wytham Tit long-term projects (detailed in the SM-S7 section). This study was supported by funding from ERC (EU, Consolidator grant 866489), FCT (Portugal, grants IF/01411/2014/CP1256/CT0007 and PTDC/BIA-EVF/5249/2014) and DST-NRF Centre of Excellence at the Fitzpatrick Institute of African Ornithology University of Cape Town awarded to RC, ANR (France, grants ANR-15-CE32-0012-02 and 19-CE02-0014-01) to CD and Marie Curie-Staff Exchange (Horizon MSCA grant 101183160) to LS, ACF, CD and RC. Blue tit data collection and AF were supported by the Canada Graduate Research Scholarship and Discovery Grant from Natural Sciences and Engineering Research Council of Canada (NSERC). Sociable weaver and blue tit long-term projects are part of the OSU-OREME, and the long-term studies in Ecology and Evolution (SEE-Life) program of the CNRS. Great tit data collection and IMB were supported by the ERC (grant 250164) and the UKRI Frontiers award EP/X024520/1 awarded to Ben C. Sheldon.

## AUTHOR CONTRIBUTIONS

LRS, ACF, CD and RC conceived the idea and methodological design. All authors contributed to their system’s video-data collection, management and analysis. LRS and ACF developed the deep-learning pipeline. LRS led the writing; all authors contributed to drafts and approved submission.

