## Supplementary Material for "From video to behaviour: an LSTM-based approach for automated nest behaviour recognition in the wild"

**Running Headline:** LSTM for behaviour identification

Liliana R. Silva ^a,b,c*^ , André C. Ferreira ^a,b,c,d^, Irene Martínez-Baquero ^e^, Arlette Fauteux ^f^, Claire Doutrelant ^d,g^ and Rita Covas ^a,b,g^

^a^ CIBIO, Centro de Investigação em Biodiversidade e Recursos Genéticos, InBIO Laboratório Associado, Campus de Vairão, Universidade do Porto, 4485-661 Vairão, Portugal

^b^ BIOPOLIS Program in Genomics, Biodiversity and Land Planning, CIBIO, Campus de Vairão, 4485-661 Vairão, Portugal

^c^ Department of Evolutionary Biology and Environmental Science, University of Zurich, Zurich 8057, Switzerland

^d^ CEFE, Univ Montpellier, CNRS, EPHE, IRD, Montpellier, France

^e^ Edward Grey Institute of Field Ornithology, Department of Biology, University of Oxford, United Kingdom

^f^ Département des Sciences Biologiques, Université du Québec à Montréal, CP-8888 Succursale Centre-ville, Montréal, Québec, Canada

g FitzPatrick Institute of African Ornithology, DST-NRF Centre of Excellence, University of Cape Town, Rondebosch 7701, South Africa

* Correspondence:

Liliana R. Silva

### SUPPLEMENTARY MATERIAL

### S1. Data collection

The sociable weaver is a colonial cooperatively breeding bird from southern Africa, living in large communal nests composed of several independent nest-chambers used for breeding. Parents are assisted by non-breeding individuals (called helpers) that help raising the young by sharing all parental care activities: provisioning, nest-chamber maintenance and nest-chamber defence. This results in variable family groups (2-12 individuals), nest-chamber visits and behaviours (from <1 to 67 visits/h). Sociable weaver nestling provisioning videos were obtained from a wild population in Kimberley, South Africa, in a relatively standardised manner from September 2014- August 2021 and were mostly filmed in full HD 1920x1080 -quality (using a Sony™ Handycam HD). Cameras are placed under the colony so that videos are filmed from a ground-up perspective, with each camera aiming to film a single nest-chamber and keeping the entrance of such at the centre of the image (see Fig. S1). This setting maintains a relatively constant recording background around the nest-chamber that is relatively similar throughout most of the videos, composed of straw agglomerations that support the nest-chamber itself and the rest of the nest structure. Each nest-chamber entrance shape might differ between nest-chambers and days (as birds continuously maintain and build around the entrance of the nest-chamber). Videos were filmed across 16 colonies, 1239 nest-chambers and 916 days, with 30 cameras, recorded mostly around early morning, but could start from sunrise to 12h after sunrise. That, together with local climatic variation, variation in background, nest-chamber structure, image quality, brightness and contrast, assured that our built models are able to be generalised across recording conditions. Video recordings were set for around two hours (2.37 ± 0.84h (mean ± SD)).

### S2. Manual behavioural analysis

Nestling provisioning behaviours through videos were manually analysed by several annotators in two steps: i) bird detection, ii) behavioural annotation and iii) individual identification. i) Bird detection was obtained through playing several video parts simultaneously in a computer screen using BSPlayer™ software, and manually adding a chapter (a timestamp saved by hitting a hotkey in BSPlayer and that can be opened later on) with the timestamp of each detected bird movement. ii) Behavioural annotation would consist on the inspection of each annotated bird movement and for each timestamp and type of behaviour i.e., time of entrance, time of exit and for each if a bird was seen building or being attacked. This effort resulted in behavioural annotation to the second of each bird detection. Finally, for each behaviour, iii) individual identification through coloured rings combination reading was assessed and annotated. This last stage was not automated as part of this work but it is associated with the manual behavioural annotation (see S6B).

### S3. Frame extraction from annotated videos

Using the manual annotations to the second, we extracted frames directly from video using “ffmpeg” (Tomar, 2006) with 1920x1080 resolution. Extracted frames from different tasks were included from distinct years, dates, starting recording hour, colonies, nest-chambers and visiting individuals.

### S4. Data pre-processing – bird detection YOLO

As behavioural manual annotations are marked to the second and not by frame (each second can have 50 frames), extracted frames from a given second still include non-interest frames with no activity (15.01 ± 4.05 frames for 44 entrances and 13.9 ± 3.33 frames for 48 exits; mean ± SD). In addition, manual annotations were not always completely synced with the moment where the behaviour actually occurred (drifting one or two seconds). This means that even by extracting frame sequences only from previously annotated timestamps we still extracted many frames of non-interest (i.e., non-activity at the focus nest-chamber). As a result, to make sure that we would extract the frames containing the behaviours of interest we extracted frames starting from one second before the timestamp of the annotation until one second after the timestamp. To filter out the excessive number of frames that contained no bird on it (i.e. the frames before and after the behaviour occurred) we trained a single-frame bird-detection model using You Only Look Once - YOLO v8 (Varghese & Sambath, 2024) from Ultralytics (<https://docs.ultralytics.com/>) to simply detect the presence of a bird in the frame (i.e., coming in and out of the nest-chamber, perching outside and sitting inside the nest-chamber; Fig.S1). For training and validation purposes, bounding boxes of birds’ location(s) in each frame were manually annotated using the LabelImg software (Tzutalin, 2018) (6107 frames for training and 766 frames for validation). We trained the medium-sized version of YOLO v8 pre-trained on COCO dataset provided by Ultralytics, for 10 epochs, with a batch size of 16 and input image size of 640x640 pixels. The remaining hyperparameters were set to the default of the Ultralytics package. This model detected bird presence with 94.78% accuracy on the validation dataset. We then implemented this model to obtain more frames to train the behaviour classification models. Then, for each manually labelled timestamp (entrance, exit, building, or aggression), we considered all frames around it in
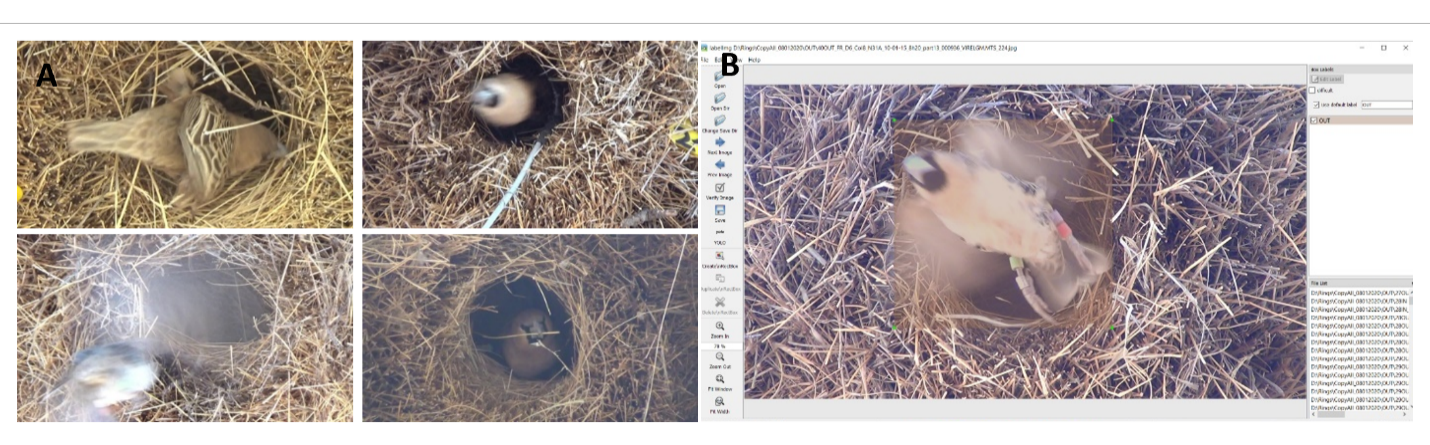
which YOLO detected a bird to belong to the same annotated behaviour.

**Figure S1**: A) Examples of frames used to train the bird detection YOLO. B) Example of a bounding box of a bird’s location(s) in a frame manually annotated using the LabelImg software (Tzutalin, 2018).

### S5. Behavioural automation

### S5A. Training and validation datasets

To train a model for detecting nest activity we used: six-frame sequences of 12443 entrance (“IN”) events, 12096 exits (”OUT”) and 15610 sequences of the negative class. For the validation dataset we used 1203 sequences of each category from videos and nests not present in training. Negative class included everything that was not a movement of interest from and to the nest-chamber including: empty nest-chamber (i.e., no visible bird on frame), perching adults, adults and chicks visible inside the nest, passing birds not moving to the focus chamber, blurred frames (e.g., from strong wind that shake the cameras and the tree), sanitation events (e.g., chicks defecating at the nest-chamber entrance), other objects used for other research purposes (e.g. microphones) and falling nest material from other chambers.

“Building” corresponds to examples where the bird is clearly seen bringing a straw while entering the nest-chamber. Negative class examples included bird entrances without bringing straw, but could bring prey. Taking advantage of having an automatic *nest activity monitoring* step, this model is applied using the already identified entrances. To detect and classify building, 6200 building and 6922 negative class six-frames sequences were used in the training dataset and 424 sequences (212 building and 212 non-building) were used in the validation dataset, from videos and nest-chambers not present in the training dataset. Additionally, if a building or non-building entrance was long enough to be divided into multiple six-frame sequences (sub-sequences of the same event), we included all of them in the training dataset. However, to obtain more accurate performance estimates and avoid data correlation, we excluded these sub-sequences from the validation dataset (i.e. each of the 424 sequences in the validation dataset correspond to an independent event).

“Aggression” examples were considered from clear observations of pecking and chasing by one or several individuals and removing the aggressed individual from the nest-chamber. Negative class examples included one bird exits with no aggression; two or more one birds exiting with no aggression and birds perched in front of the nest-chamber. Taking advantage of having an automatic *nest activity monitoring* step, this model was applied using the already identified exits. To detect and classify this behaviour, 633 aggression and 891 negative class frames were used for the training dataset and 192 (96 aggressions and 96 non-aggressions) were used for the validation dataset. Similarly to the building dataset and given the limited data to train and evaluate the models, we selected aggression (and non-aggression) instances for the validation dataset from videos and nest-chambers not present in the training dataset. Furthermore, similarly to the building models only unique sequences of aggression (and non-aggression) were included in the validation dataset (i.e. all instances come from independent events). Since it typically takes at least 16 frames for humans to recognise aggressive behaviour, we created six-frame sequences skipping two frames in between. This means that each sequence was constructed from an original set of 16 frames, where we kept only one frame for every three.

### S5B. Behavioural classification with recurrent network(s) – LSTM(s)

After pre-processing, the sequenced data was framed into windows of six frames to train long short-term memory networks (LSTM). We developed separate models for nest activity, building and aggression behaviours to take advantage that building and aggression are associated with specific nest activity movements (building only happens during a nest entrance and aggression always results in an exit) and, therefore, using multiple models allows to implement a hierarchal approach in which the building and aggression models are only run if a IN or OUT, respectively, are detected. For the nest activity model, we used a VGG19-backbone pre-trained on ImageNet for which the final classification was removed. The VGG19 layers were wrapped in a “TimeDistributed” layer to process each frame independently. A global max pooling 2D layer was added to reduce spatial dimensions and was followed by an LSTM layer with 1024 units to capture temporal dependencies across the frame sequence. After the LSTM layer, we added four dense layers with 1024, 512, 128 and 64 units and ReLU activation functions to reduce the feature space while learning non-linear combinations of the features for classification, which was done by adding a final layer with a softmax activation function and three units corresponding to each of the categories: IN, OUT and negative class. For the other two models (aggression and building classification) a one neuron layer was added with a sigmoid activation for classification (since there were only two classes: building or not building; aggression or no aggression, for the building and aggression models, respectively). The categorical cross-entropy was used as the loss function for the nest activity model and binary cross-entropy for the other two models. For the aggression and building models, instead of using a VGG19 back-bone pre-trained on ImageNet and layers with random weights initialisation we used the model trained for nest activity as a starting point, only modifying the last classification layer (from three units and a softmax function to one unit with a sigmoid function). For all models we used SGD optimiser with a learning rate of 1e-3 for all models and a batch size of one (i.e. a sequence of six images). Models were trained using an RTX 3090 GPU.

For training the aggression and building models, given the relatively small and imbalanced dataset, we under-sampled the majority class in each epoch to ensure the model was equally exposed to both positive and negative classes. Furthermore, for these two models we applied data augmentation to prevent overfitting. We used similar image transformations as in Ferreira et al., 2020, which included Gaussian blur, motion blur, cutout (randomly masking out square regions of the image). These transformations were randomly applied alone or in combination.

The nest activity model was trained until there was no improvement in the loss for more than one epoch. The other two models were trained until there was no more improvement in the loss for more than 10 consecutive epochs. The trained models allowed for a classification in video at the selected frame-interval, outputted in a categorical class, translated to a 1s output time data classification (our meaningful biological resolution of analysis) and an associated prediction accuracy that allows to pre-filter classifications of lower confidence (i.e., filter false detections – avoid false positives (“FP”), but not exclude true detections – avoid false negatives (“FN”). For all models we only considered a predicted behaviour to be true if the output values’ confidence was above 0.9. Otherwise, the predictions were considered as “negative class”. This was done to prevent a large production of false detections and the threshold was chosen before running the models on the testing data (see below).

### S6. Evaluation

For evaluating the models under a real application scenario, we used complete videos that were not included either in the training or the validation datasets. For the nest activity model, we used 315 videos, for the building model we used 69 videos and for the aggression model, we used 23 videos (sample sizes vary because not all videos contained building and aggression events as they are low frequency events). Videos were processed using the CPU version of “decord” package in python (<https://github.com/dmlc/decord>). Videos were broken into non-overlapping sequences of six frames. All sequences were processed by the nest activity model first to predict: IN, OUT or negative class (i.e. no activity). If an IN event was detected, we then passed the six-frame sequence to the building model to predict building or non-building. If an OUT event was detected, then a new 16-frame sequence was generated by including the five frames before the OUT event, the six frames of the detected-OUT event and the following five frames. The sequence was then compacted into a six-frame sequence by only keeping one frame every three frames. The resulting window was then passed to the aggression model to predict aggression or non-aggression.

All models were evaluated against human observers’ annotations, which we considered as representing ground-truth annotations. Since our behavioural segments are based on human annotations, they are inherently prone to error and subjectivity. Therefore, we have employed several strategies to minimize such labelling errors. First, validation videos were screened extensively: each video was reviewed twice by two independent annotators, with the second observer always being experienced, to establish ground-truth labels for each video. Additionally, logical consistency checks were used during correction to identify behaviourally impossible patterns (e.g., bird exits without prior entries, attack events without the presence of two individuals). Importantly, this was also done even if the first analyser was experienced, as we did not assume that experienced observers are error-free. We explicitly quantified annotation error across both experienced and non-experienced observers based on annotators’ disagreement (after carefully examining all disagreements to understand which annotation was correct), and report these results in the Table S1, where observer performance is discriminated by experience level and observer identity. Because the sociable weaver dataset is part of a long-term project involving multiple annotators over several years, we used the average human annotation error across all observers, independent of experience level, as the benchmark for comparison with model performance (reported in Table S1). In addition, it is important to note that all “unexperienced” observers go thought an intensive training process of at least one month before their data is used. We report observers’ performance only as false positives and false negatives because observers were consistently accurate in identifying the type of behaviour (e.g., correctly labelling an entrance as an entrance). Observer errors’ primarily stem from missing moments of activity and from inaccuracies in noting the precise time point at which a behaviour occurs. For example, they may miss a bird entering the nest while watching the video, resulting in a false negative. Conversely, they might mistakenly annotate irrelevant events, such as falling nest material that resembles a bird exiting, as false positives. We believe this provides a complete and realistic estimate of human error during behavioural annotation. Secondly, further quality control was applied at the frame extraction stage. Training data were also screened prior to model fitting: extracted frames from all training were manually reviewed. This implied the visual inspection of all frames in a given frame interval (previously annotated for a given behaviour) to access if their labelling was correct.

Finally, the biological significance of our behavioural classifications was confirmed by assessing if our automated data was able to capture meaningful trends confirmed in our long-term study (see below for each behavioural task).

Moreover, by extensively screening our training and validation data for potential inaccuracies, we believe the remaining error rate is minimal. Nonetheless, given the large number of annotated events, some errors may have gone undetected despite our careful screening. However, the models we developed have demonstrated strong performance and generalisation, indicating robustness to occasional annotation noise (also a well-known a property of large datasets, Rolnick et al., 2017). This resilience suggests that the overall patterns in the data are sufficiently consistent to support reliable model training, even in the presence of minor inaccuracies, and adequate comparison to humans’ performance.

Comparison between model performance and speed to annotators, as well as biological significance, were conducted using R v. 4.2.3 (The R Foundation for Statistical Computing, Vienna, Austria, <http://www.r-project.org>). All model and human performance analyses below were conducted using the same software.

### S6A. Nest activity monitoring evaluation

Using our observers’ manual analysis of movements around the nest-chamber as possible entrances and exits (described in S2), we were able to compare manual annotations of different observers, experienced (> six months of analysis experience) and non-experienced, with the model detections to assess the introduced error (Table S1). Annotations and detections were compared in terms of the proportion of false positives (i.e., when a moment is annotated and no bird is entering or exiting the nest-chamber) and false negatives produced (i.e., when a bird is entering or exiting the nest-chamber and no moment was annotated). These errors were detected by reviewing these videos a second time by a different annotator. Nonetheless, this error value is underestimated as manual annotation encompasses the manual writing of timestamps and behaviour, which introduces additional errors not accounted for here, whereas the model outputs data automatically.

**Table S1:** Comparison of model and human performance in detecting sociable weaver nest activity (i.e., detecting nest-chamber entrances and exits) using the percentage of false positives (“FP”) and false negatives (“FN”) (i.e. number of false positives or negatives/number of visits). False annotations were averaged across all videos for each annotator. For the manual analysis, we randomly selected 20 videos for each of the six observers (total of 244h and 6,568 movements), distinguishing between experienced (> six months analysing) and non-experienced (< six months analysing). For model analysis, we randomly selected 100 videos (total of 210h and 6,079 movements) to be processed by the model**.**

| **Observer** | **FP (%)** | **FN (%)** | **Videos (nbr)** |
| --- | --- | --- | --- |
| Experienced1 | 4.08 | 0.59 | 20 |
| Experienced2 | 3.89 | 0.8 | 20 |
| Non-experienced1 | 8.97 | 5.97 | 20 |
| Non-experienced2 | 20.31 | 10.05 | 20 |
| Non-experienced3 | 13.11 | 1.44 | 20 |
| Non-experienced4 | 12.41 | 1.33 | 20 |
| Average observers | 10.05 | 3.36 | 120 |
| **Model (Nest activity)** | 6.24 | 0.39 | 100 |

To assess speed, we compared the time taken by five observers, both experienced and non-experienced, and by the model to mark the timestamps of nest activity in 2hs videos (Table S2). Nonetheless, this speed value is underestimated as our model simultaneously performs detection and behaviour classification (distinguishing entrances from exits), and automatically outputs data frames with classifications and associated times, while annotators at this stage are simply marking timestamps without behaviour classification.

**Table S2:** Comparison of model and human analysis speed (minutes). For manual analysis, we randomly selected videos (of 2hs) for five observers (70 videos), differentiated as experienced (> six months analysing) and non-experienced (< six months analysing). For model analysis, we randomly selected 25 videos (of 2hs).

| **Observer** | **Total speed (min)** | **Videos (nbr)** |
| --- | --- | --- |
| Experienced | 49.8 | 19 |
| Non-experienced1 | 51.00 | 12 |
| Non-experienced2 | 38.55 | 9 |
| Non-experienced3 | 33.85 | 13 |
| Non-experienced4 | 29.29 | 17 |
| Average observers | 40.5 | 70 |
| **Model (Nest activity)** | 28.6 | 25 |

To biologically validate our activity detection model, we assessed whether our automated movement detections produced data consistent with a known biological pattern in the sociable weaver nestling provisioning stage: an increase in nest-chamber visits with increasing nestling age and number of nestlings, reflecting higher feeding demand. To understand this visiting pattern in our manual data, we applied a generalised linear model, using the “lmer” function from the “lme4” package in R (Bates et al., 2015) to 1497 manually analysed nestling provisioning videos. Total visits to the nest were fitted as the dependent variable, with three independent variables: nestling age (in days), number of nestlings and total video length (in minutes). Additionally, laying date, colony and season were fitted as random variables. We applied the same analysis to 100 automatically analysed nestling provisioning videos. Due to singularity issues, we were not able to fit any random variable, and therefore we applied a linear model, using the “lm” R core function (Bates et al., 2015) with the same dependent variables mentioned above. The number of detected entrances was used as a proxy for the number of visits. To verify the normality assumptions, the models’ residuals were analysed using QQ plots, fitted versus residuals plots and histograms. All predictors were scaled to allow direct comparison of estimates. Both manual and automated analyses showed the same significant positive relationship (Fig. S2): an increase in the number of visits with nestling age (β=7.70 ± 0.45, t=17.15, P < 0.01 and β=8.24 ± 1.76, t=4.67, P < 0.01, respectively) and with the number of nestlings to be fed (β=6.68 ± 0.57, t=11.66, P < 0.01 and β=5.38 ± 1.76, t=3.05, P < 0.01, respectively).

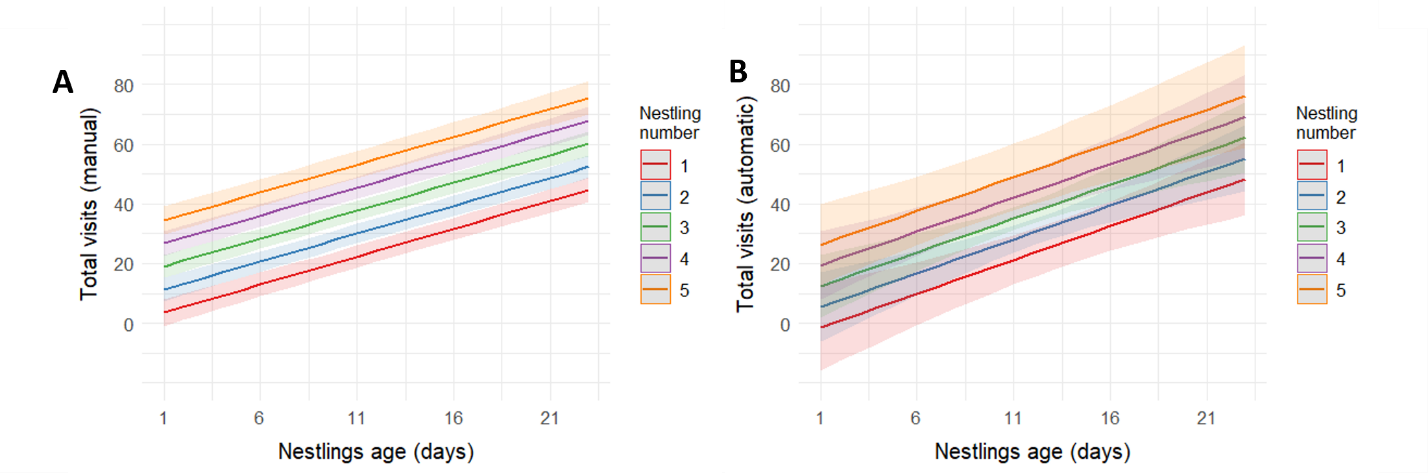
**Figure S2:** Predicted total visits to the nest with increasing nestling age, number of nestlings in the nest and video length for (A) manual (n=1,497 videos with 54,609 visits) and (B) model annotations (n=100 videos with 3,405 visits), using linear models. Solid lines represent model predictions, colours represent nestling count category, and coloured shadows represent model confidence intervals.

### S6B. Building and aggression detection evaluation

Following the same procedures as above, we were able to compare building and aggression manual annotations by experienced and non-experienced observers with the model detections to assess error in terms of false positives and negatives (Table S3 and S4). The number of observers differs between activities, as some were not instructed to annotate some specific behaviours (e.g., for building, only one of the five non-experienced observers systematically annotated this behaviour).

**Table S3:** Comparison of model and human performance in detecting sociable weaver building behaviour (i.e., if a bird is entering with a straw), as the percentage of false positives (“FP”) and false negatives (“FN”) (i.e., number of false positives or negatives/number of visits). False annotations were averaged across all videos for each annotator. We used 69 videos (of around 2hs) with 339 building events for a total of total of 2,004 entrances, analysed by two observers-one experienced (> six months analysing) and one non-experienced (< six months analysing), and further processed by the model.

| **Observer** | **FP (%)** | **FN (%)** | **Videos (nbr)** |
| --- | --- | --- | --- |
| Experienced | 0.8 | 1.15 | 34 |
| Non-experienced | 0.36 | 8.8 | 35 |
| Average observers | 0.58 | 4.97 | 69 |
| **Model (Building)** | 3.97 | 2.3 | 69 |

**Table S4:** Comparison of model and human performance in detecting sociable weaver aggression behaviour (i.e., if a bird is attacked out of the nest-chamber) as percentage of false positives (“FP”) and false negatives (“FN”) production (i.e. number of false positives or negatives/number of visits). False annotations were averaged for all videos for each annotator. We used 23 videos (of around 2hs) with 35 aggression events for 1,061 exits, analysed by four observers differentiated as experienced (> six months analysing) and non-experienced (< six months analysing), and further processed by the model. As few videos were analysed by three different “non-experienced” annotators, their respective results were compiled and are presented together.

| **Observer** | **FP (%)** | **FN (%)** | **Videos (nbr)** |
| --- | --- | --- | --- |
| Experienced | 0.11 | 0 | 12 |
| Non-experienced | 3.13 | 6.44 | 11 |
| Average observers | 1.62 | 3.22 | 23 |
| **Model (Aggression)** | 5.82 | 0.06 | 23 |

Regarding speed evaluation, we could not isolate the time required for manual behavioural detection (i.e., identifying building or aggression), as observers assessed behaviours only after annotating nest-chamber movements, going through these annotated movements, identified other behaviours (i.e., if building or aggressing), together with individual identification (via colour band reading, which also required time). Therefore, we compared the model speed with the time taken for manual behavioural annotation (of both building and aggression) plus individual identification, which could take an experienced annotator 17.73 ± 10.26 min (mean ± SD, n= 41 videos). The model speed for detecting building and aggression behaviours was less than 1 min per video.

To biologically validate our building detection model, we assessed if our automated building detections would show a similar biological difference between incubation and nestlings’ nest stage: during incubation, although with much fewer visits, building is seen more often in sociable weavers. To describe this building pattern in our manual data, we applied a binomial logistic regression model, using the “glmer” function from the lme4 package, to predict the proportion of building visits out of total visits, based on the nest stage for 752 manually annotated videos. The independent variable was modelled as the number of building visits to the non-building visits (total of visits minus the building visits). Nest-chamber stage was included in the model as a dependent variable and was defined as having “eggs” or “nestlings” in the nest. Due to singularity issues, only the brood ID, was fitted as a random variable. In resemblance, we assessed the same relationship to 72 automatically analysed videos. Again, due to singularity issues, we were not able to fit any random variable, and therefore we applied a generalised linear model with the same configuration as above but without random terms. The number of building detections was used as a proxy for the number of building attempts. Both manual and automated videos showed the same significant positive relationship of increased building events for incubation (Fig. S3): β=-2.97 ± 0.21, t=-14.35, P < 0.01 and β=-0.45 ± 0.18, t=-2.44, P = 0.01, respectively.

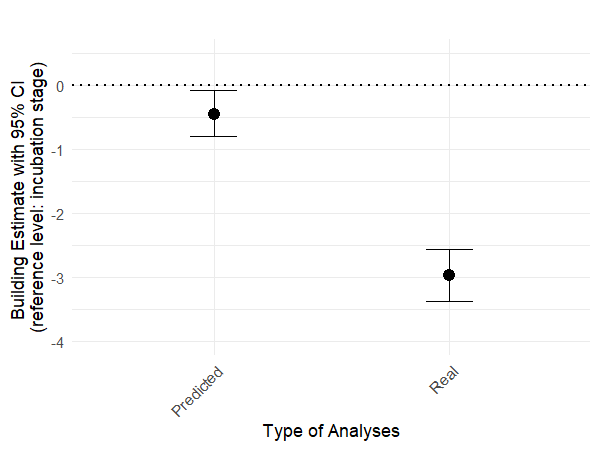
**Figure S3:** Estimated effect sizes (log-odds) with 95% confidence intervals (CI) from the GLM predicting the likelihood of building (yes/no) per visit, as a function of breeding stage (incubation vs. nestling). Predicted and real likelihood of building correspond to model (72 videos, 5479 visits, of which 287 building events) or human annotations (756 videos, 21634 visits, of which 1,066 building events), respectively. Negative estimates with CIs that do not overlap zero indicate a reduced likelihood of building during the nestling stage. Modelling details in Supplementary Material S6.

For aggression, as an extremely low frequency behaviour observed and because we did not have any clear a priori prediction regarding this behaviour, we did not test for any biological pattern in the sociable weaver.

### S6C. Pipeline speed evaluation

For the speed evaluation, we took into account that computers can operate continuously 24/7, while annotators can work about eight hours a day, five days a week. Based on the speed values reported above, we estimated that the computer could process 345.68 two-hour videos per week, compared to 41.25 two-hour videos per week for humans.

### S6D. Additional metrics

**Table S5:** For complete model evaluation, the Macro F1 score is reported for each sociable weaver model. This metric represents the harmonic mean of precision and recall, computed per class and averaged equally across classes, with values closer to 1 indicating better performance.

^a^Nest-chamber entrance, ^b^nest-chamber exit, ^c^negative class

| *Model* | Classes | Macro F1 score |
| --- | --- | --- |
| *Nest activity* | IN^a^ vs. OUT^b^ vs. NC^c^ | 0.93 |
| *Building* | building  vs. NC^c^ | 0.88 |
| *Aggression* | aggression  vs. NC^c^ | 0.88 |

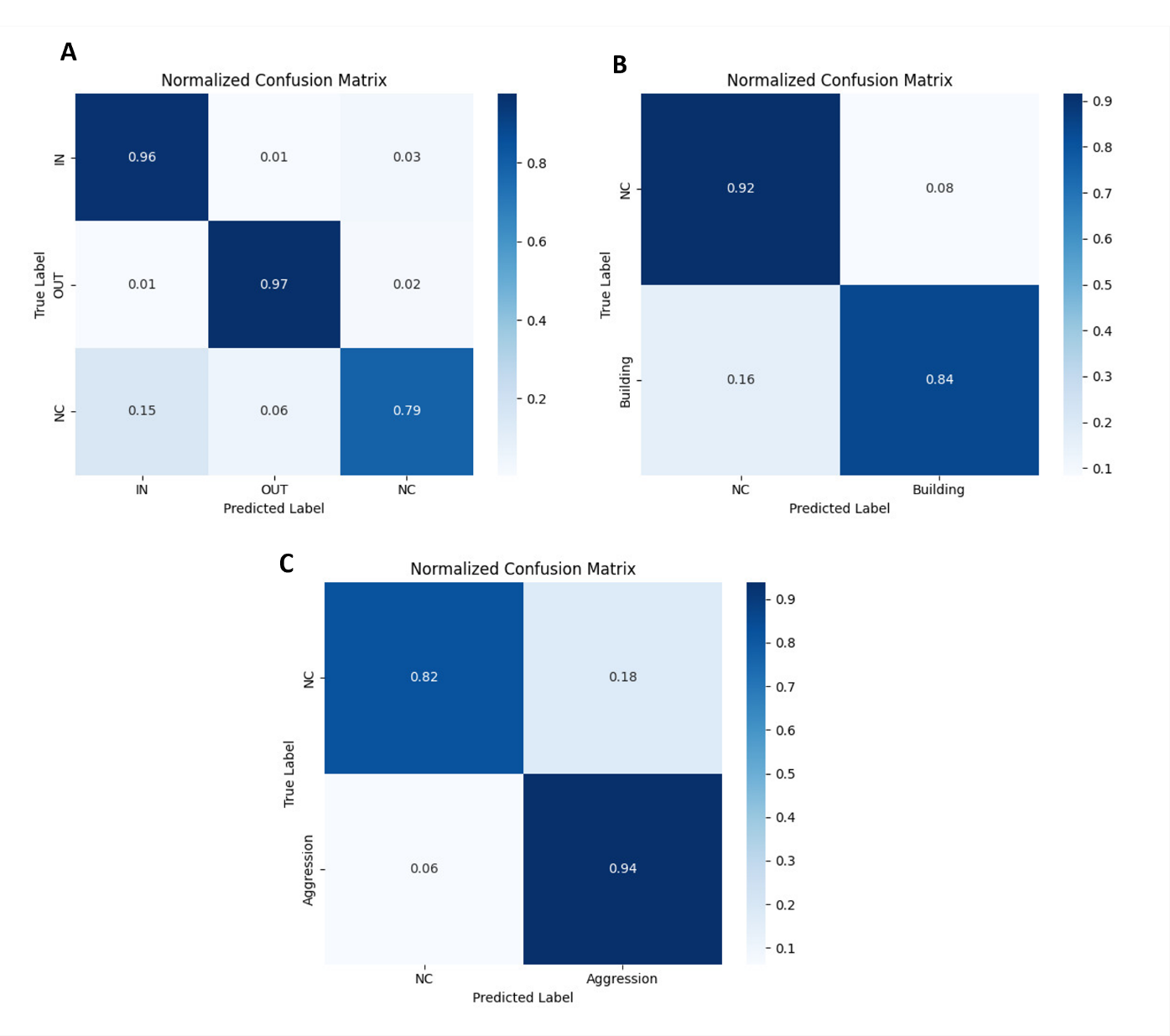
**Figure S4:** Normalized confusion matrices are presented for each sociable weaver model to illustrate classification performance for a) nest activity, b) building and c) aggression detection. These matrices display the proportion of predicted versus true classes, highlighting both correct classifications and misclassifications, and complement the Macro F1 evaluation by showing class-level performance.

**C**

### S6E. Importance of hard negatives

We tested the effect of the percentage of hard negatives present in the negative class on the nest activity dataset. To test this, we built two datasets of 11,146 sequences. In one dataset, we included only 10% of hard negatives in the “NC” category. This proportion of “NC” reflects approximately the proportion we would find if we randomly sampled the videos and selected timestamps without any INs, OUTs, building or aggressions. The other dataset contained 55% of hard negatives and represents our effort to target hard “NC” instances. From each of the datasets, we sampled 1,113 sequences for validation, and the remaining sequences were used for training (10,033). Sequences used for validation were obtained from different videos and nests than the ones used in the training datasets. In addition to the sequences for validation (used to compute “Model accuracy % (sequences)” in Table S6), we also tested the models on 33 videos (used to estimate “Model FP% | FN% (videos)” in Table S6). We trained both models following the same procedure as described above (S5B). Results are shown in Table S6.

In both models (“10% hard negatives” and “55% hard negatives”), the proportion of hard negatives was kept the same in the validation and training datasets (i.e., for the “10% hard negatives” model, there are 10% hard negatives in the training and in the validation dataset, the later dataset was used to estimate the model accuracy named “Model accuracy % (sequences)”, mentioned hereafter as “accuracy”). The slight differences in accuracy can be attributed to either i) the fact that the “55% hard negatives” validation dataset has more hard negatives and, therefore, more challenging to classify examples, or ii) just a result of model training stochasticity since we only trained one model for each scenario. However, our main point of these analyses was to demonstrate that low quality validation (and training) datasets (here the “10% hard negatives” dataset) can generate misleading metrics. This is highlighted by the fact that, even though the “10% hard negatives” dataset generated slightly better accuracy, there are marked differences during deployment, i.e., “Model FP% | FN% (videos)”, with more than double the false positives production for the “10% hard negatives” dataset; when both models were evaluated on the same real-world videos. This exercise was designed to demonstrate that when training and validation datasets lack sufficient hard negatives, models may appear to perform well during training and validation (e.g. exhibiting higher accuracy), yet perform poorly during deployment because they were not adequately exposed to the challenging examples that most strongly affect real-world application.

**Table S6:** Training details and performance of sociable weaver nest activity models with varying percentages of hard negatives. Training details include: classes, interval of frames, training and validation datasets sizes. Examples of negative class (“NC”) allow to include variation in the models. Performance was assessed as the model frame-accuracy obtained at the validation stage, but also, by assessing the produced false positives and false negatives, performed by the models for 2h videos.

^a^Nest-chamber entrance, ^b^nest-chamber exit.

| *Model* | Classes | Frames  training | Sequences  validation | Model accuracy %  (sequences) | | Model  FP% \| FN%  (videos) |
| --- | --- | --- | --- | --- | --- | --- |
| *10% hard negatives* | IN^a^ vs. OUT^b^ vs. NC | 10,033 | 1,113 | 97.30 | 19.00 \| 1.02 | |
| *55% hard negatives* | IN^a^ vs. OUT^b^ vs. NC | 10,033 | 1,113 | 94.44 | 7.67 \| 0.96 | |

### S6F. Effect of training dataset size

To test the effect of the sample size of the training data we trained different models for nest activity that varied on the amount of data used during training. From the initial dataset of 43,758 sequences, we selected 1,203 sequences from each category (IN, OUT and NC) for validation and the remaining data was used for the varying training dataset sizes (25% of the data, 50%, 75% and 100%, respectively, Table S7). Sequences used for validation were from different videos and nests than the ones used in the training datasets and were the same for all models. In addition, to using the validation sequences, we also evaluated the models on 31 videos. We trained all models following the same procedure as described above (section S5B). Results are shown on Table S7. Models were overall similar in their good performance, independently of the dataset size, with differences in error so low that can be attributed to model training stochastic. The absence of consistent performance gains with increasing dataset size is likely attributable to the well-established phenomenon of diminishing returns with respect to training data in deep learning. In many machine learning settings, model performance improves rapidly when moving from very small to moderate sample sizes. However, beyond a certain point, learning curves tend to plateau, such that further increases in training data yield no, or only marginal improvements in performance, (see for example, Norouzzadeh et al., 2018). In our case, performance appears to have plateaued at comparatively small sample sizes (already at 25% of the full dataset), but reducing it further starts to substantially decrease performance (e.g., at 12% of the full dataset, the FP raises to 7.40% and FN to 6.84%, see section S6G below). One possible explanation is that by deliberately incorporating hard-to-classify examples (e.g., hard negatives) into the training set, thereby increasing sample diversity, we have improved the coverage of decision-boundary regions of all our different models. Because difficult examples disproportionately influence gradient updates, their inclusion may have enabled the models to learn robust decision boundaries, even with reduced training data.

**Table S7:** Training details and performance of sociable weavers’ nest activity models of different sizes. Training details include: classes, interval of frames, training and validation datasets sizes. Examples of negative class (“NC”) allow to include variation in the models. Performance was assessed as the model frame-accuracy obtained at the validation stage, but also, by assessing the produced false positives and false negatives, performed by the models for 2h videos.

^a^Nest-chamber entrance, ^b^nest-chamber exit.

| *Model* | Classes | Sequences  training | Sequences  validation | Model accuracy %  (sequences) | | Model  FP% \| FN%  (videos) |
| --- | --- | --- | --- | --- | --- | --- |
| *25% data* | IN^a^ vs. OUT^b^ vs. NC | 10,037 | 3609 | 95.32 | 2.64 \| 4.63 | |
| *50% data* | IN^a^ vs. OUT^b^ vs. NC | 20,074 | 3609 | 93.82 | 3.85 \| 0.80 | |
| *75% data* | IN^a^ vs. OUT^b^ vs. NC | 30,112 | 3609 | 95.76 | 1.81 \| 3.48 | |
| *100% data* | IN^a^ vs. OUT^b^ vs. NC | 40,149 | 3609 | 92.02 | 4.63 \| 1.39 | |

### S6G. LSTM vs YOLO

YOLO, due to its easy implementation, is often the first approach attempted by researchers when solving an image-based classification problem, including for behavioural analyses. Although YOLO is not designed for action recognition, a recent study (Chan et al., 2025) demonstrates that it can perform competitively and even outperform another commonly used framework for automated video analyses (e.g., DeepEthogram), including for nest visiting behaviour. Moreover, to compare this single frame approach with the LSTM approach we built two training datasets, one containing 4,956 sequences (corresponding to ca. 12% of our full dataset) of 6 frames to train the LSTM and another dataset containing 4,956 images manually annotated with the delimitation of a bounding around the bird and its behaviour (“IN” or “OUT”, see Fig. S1 for more details) or a bounding box around an object or bird that could be mistaken by a bird going in or out (i.e. “NC”; nestlings, material falling off the nest, bird passing in front of the camera and other instances). Although the LSTM model used six times more frames (as each sequence contains six frames), we consider this a fair comparison. First, these correspond to the natural ‘statistical units’ of each model (a sequence for LSTM, a single image for YOLO). Second, while sequences involve more frames, the within-sequence variation is limited (e.g., same nest, background, and individual bird). Third, and most importantly, annotating 4,956 frames with bounding boxes for YOLO required considerably more time than extracting 4,956 sequences for LSTM, since the latter only needed to be linked to existing annotations from already manually annotated videos and extracted in groups, without additional manual effort. In addition, 1500 frames have been suggested to be enough to train models for behaviour classification using YOLO (Chan et al., 2025).

To train the YOLO models we selected 540 pictures for validation (180 per category). We trained the version of YOLOv8 for 150 epochs, with the default settings of Ultralytics (2023). The LSTM model was trained similarly by selecting 540 sequences for validation and we followed the same training procedures as described above in S5. For YOLO, accuracy was estimated on the validation dataset by considering a correct prediction if the bounding box with the highest confidence overlapped with the ground truth bounding box on an IoU threshold of 0.05 and predicted the correct label.

In addition to the validation frame-based dataset, both models were evaluated on 31 videos as in S6A. For both models, we selected 7 videos to define a confidence threshold to exclude soporiferous predictions (threshold used on the label confidence for YOLO was 0.7, LSTM was kept the same optimal 0.9). For YOLO, since predictions are per frame, after applying the threshold we grouped together frames within one second with the same prediction as belonging to the same behaviour event. A similar rule was applied for the LSTM, two sequences within the same second with the same prediction would be considered as the same behaviour event. LSTM largely surpassed both types of YOLO models (Table S8).

**Table S8:** Training details and performance of sociable weavers’ nest activity of YOLO and LSTM modelling approaches. Training details include: classes, interval of frames, training and validation datasets sizes. Examples of negative class (“NC”) allow to include variation in the models. Performance was assessed as the model frame-accuracy obtained at the validation stage, but also, by assessing the produced false positives and false negatives, performed by the models on 2h-videos.

^a^Nest-chamber entrance, ^b^nest-chamber exit, ^c^negative class

| *Model* | Classes | Frames  interval | Frames or sequences  training | Frames or sequences  validation | Model accuracy %  (sequences) | Model  FP% \| FN%  (videos) |
| --- | --- | --- | --- | --- | --- | --- |
| *YOLOv8* | IN^a^ vs. OUT^b^ vs. NC^c^ | - | 4,956 | 540 | 80.56 | 54.70 \| 1.06 |
| *LSTM* | IN^a^ vs. OUT^b^ vs. NC^c^ | 6 | 4,956 | 540 | 96.11 | 7.40 \| 6.84 |

### S7. Blue tit behavioural automation

### S7A. Blue tit data collection and manual behavioural analysis

Blue tits (*Cyanistes caeruleus*) nest-boxes were recorded between April and June 2024, in Corsica (France), to investigate the link between the provisioning and exploratory behaviour of blue tit adult birds. As the quality and quantity of prey in a nestling’s diet significantly impact their development (Blondel et al., 1991), exploratory behaviour, through its link to prey choice, could be associated with reproductive success.

Videos were recorded with HQ using Wireless Bird Box Cameras^©^ (Green-Backyard). Cameras were installed through a hole located on the side of the nestboxes (modified from the wooden nestboxes of LPO collection) and were protected by a special plastic box. Recordings started when nestlings were 8 days old until they were approximately 18 days old. Recording was continuous (24h) except when batteries were changed and were emptied. The videos were stored directly in the cameras on a microSD card and recorded in blocks of 2 hours. Two populations were included in the study, both situated in the north of Corsica (Muro-D: 42°32′N, 08°55′E, 350m of elevation; and Pirio-E: 42°34′N, 08°44′E, 200m of elevation), totaling 17 nestboxes. In total, the recording effort resulted in over 4,000 hours of footage, yielding 185 videos.
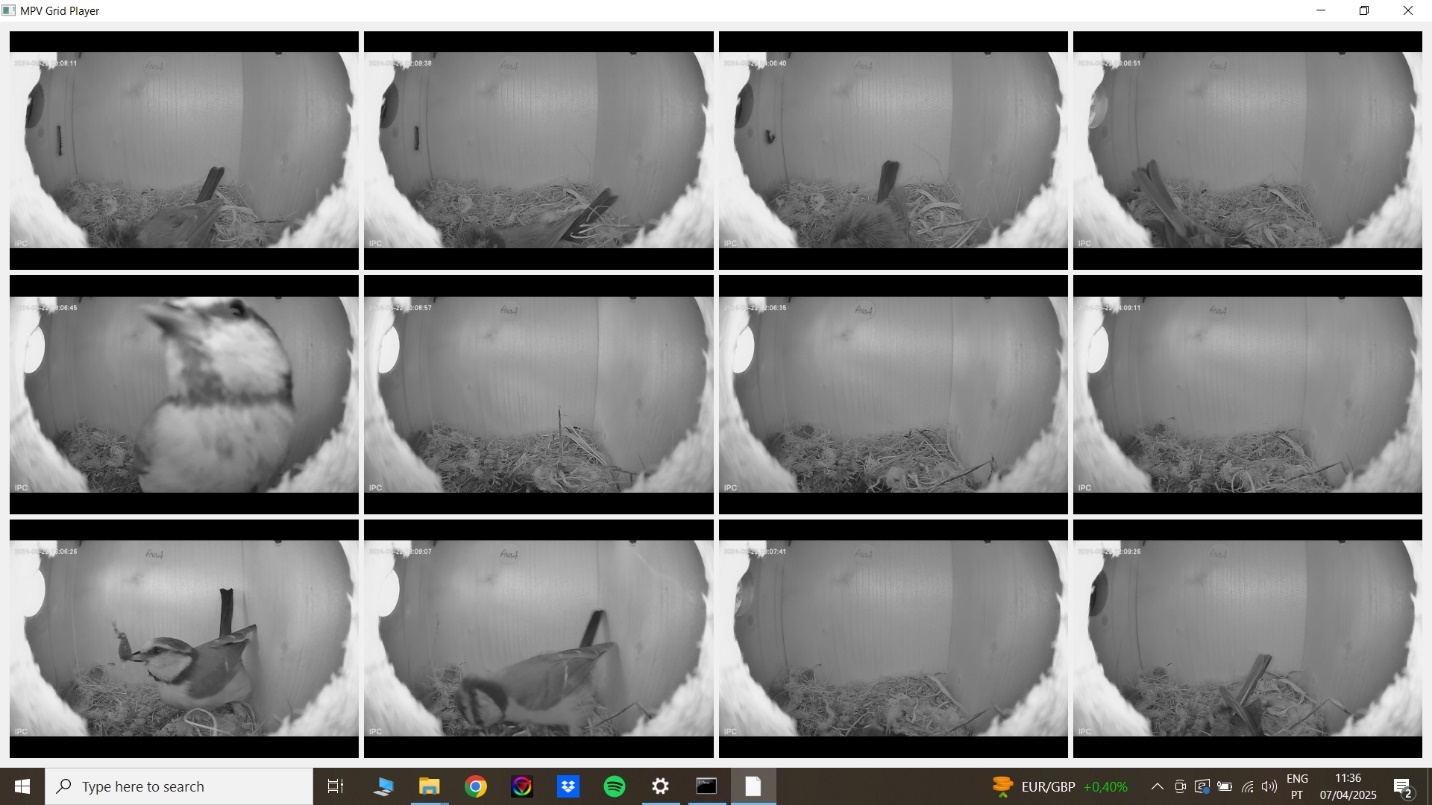
Since the sampling effort was greater and involved longer videos, conducting a behavioural analysis of each full-length blue tit video could become too time-consuming. Moreover, since the 24-hour videos were saved in 2-hour segments, behavioural annotation was performed using a custom-built analysis interface that allowed simultaneous viewing of 12 sampled segments from different nests and videos at double speed (Fig. S5). Entrances (“INs”) and exits (“OUTs”) where recorded in similarity to the great tits (see below), but we recorded additionally when sanitation was observed (i.e., when a bird picks and carries a faecal bag to outside of the chamber, “sanitation”). To encompass all variation, without requiring to annotate the full videos, we have sampled each day of each nest recording, for at least one part of 2hs, encompassing all different times in the day of recording between nests.

**Figure S5:** Custom Python-based graphical interface for simultaneous behavioural annotation of multiple blue tit nest-box videos for a streamline analysis. Built using PyQt5 and the MPV video backend, the tool allows to view and annotate up to 12 video segments at once, played at user-defined speeds (e.g., 2×). Left- and right mouse-clicks register behavioural events (e.g., "IN" or "OUT", respectively), which are saved with precise timestamps to a CSV file.

### S7B. Blue tit frame extraction from annotated videos

The same custom frame extraction and labelling interface as the great tits (see below) was used for the blue tit data, with two significant changes. The first, because the frame rate was lower, every annotation was screened in a 6 frames basis. Second, in addition to entrances and exits, we annotated “sanitation” events.

### S7C. Blue tit LSTM modelling

We trained an LSTM model for blue tits following the same procedures described in S5B. Sanitation identification was performed using a hierarchical approach similar to that applied for aggressions and building in the sociable weavers, by detecting it in a separate model of exits.

### S7D. Blue tit LSTM evaluation

Similar to the approach applied for the sociable weaver automation, we evaluated our blue tit models under a real application scenario. The evaluation was based on 18 two-hour videos from nine different nests, totalling 36 hours of footage and comprising 1210 nest movements and 160 sanitation events. These videos were manually analysed and corrected to establish a ground truth dataset, covering the widest possible range of filming days and times. The models for nest activity (“IN” and “OUT” events) and sanitation automatic detection were then applied to these videos as described in Section “S6. Evaluation”, to assess error rates and processing speed only, since the dataset was too limited to allow a direct comparison with manual annotators. Based on the preliminary analyses of 3 videos we considered a predicted behaviour to be true if the output values confidence was above 0.6.

### S7E. Blue tit LSTM additional metrics

**Table S9:** For complete model evaluation, the Macro F1 score is reported for each blue tit model. This metric represents the harmonic mean of precision and recall, computed per class and averaged equally across classes, with values closer to 1 indicating better performance.

^a^Nest-chamber entrance, ^b^nest-chamber exit, ^c^negative class

| *Model* | Classes | Macro F1 score |
| --- | --- | --- |
| *Nest activity* | IN^a^ vs. OUT^b^ vs. NC^c^ | 0.99 |
| *Sanitation* | sanitation  vs. NC^c^ | 0.93 |

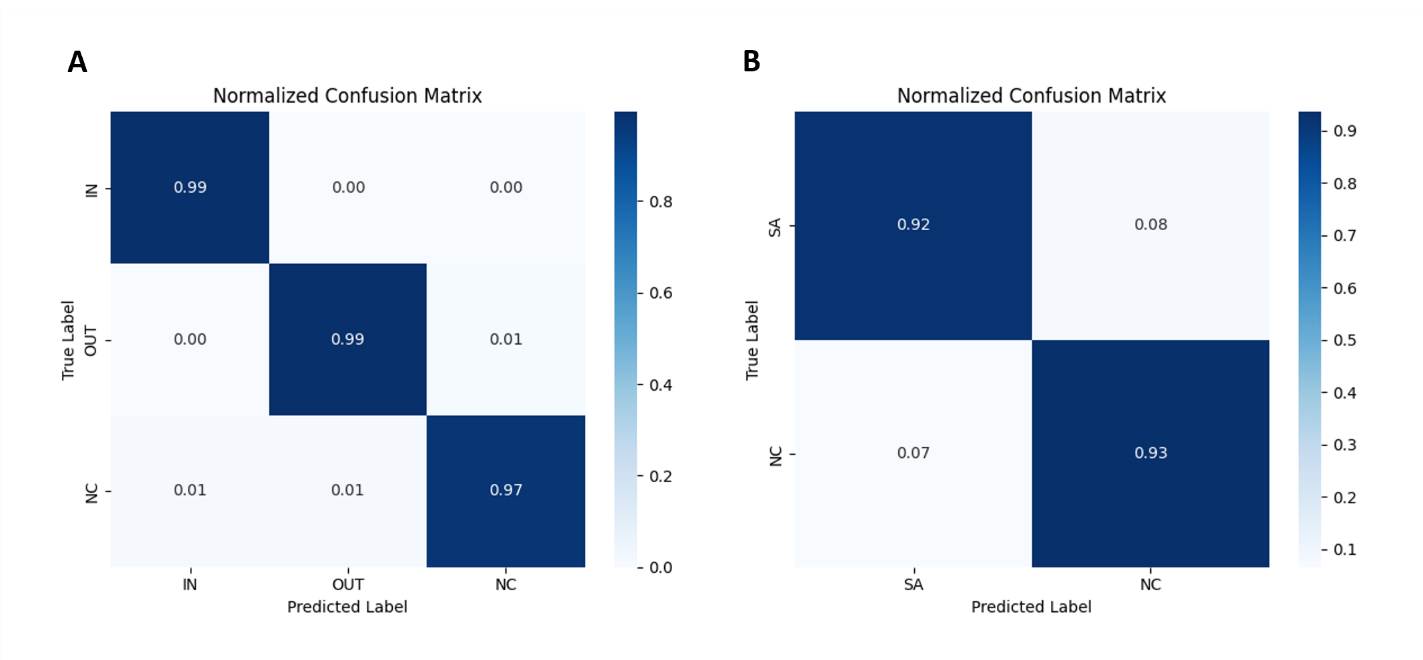

**Figure S6:** Normalized confusion matrices are presented for each blue tit model to illustrate classification performance for a) nest activity and b) sanitation detection. These matrices display the proportion of predicted versus true classes, highlighting both correct classifications and misclassifications, and complement the Macro F1 evaluation by showing class-level performance. Matrix A and B correspond to the nest activity and sanitation detection models, respectively.

### S8. Great tit behavioural automation

### S8A. Great tit data collection and manual behavioural analysis

Nest-box breeding bird populations provide an opportunity to investigate how birds adjust foraging in phenologically heterogeneous landscapes. In such systems, spatio-temporal mismatches between peak prey availability and chick demands create a strong selective pressure on parental provisioning strategies (Hinks et al., 2015; Wilkin et al., 2009). By extracting behavioural traits from video footage, such as visit rate, we can estimate optimal foraging predictions under shifting trophic synchrony.

In May 2024, we recorded nest provisioning events in a wild population of nest-box-breeding Great tits (*Parus major*) in Wytham Woods (Oxfordshire, UK). We used Camera Module V3 NoIR units with 120° wide-angle lenses placed inside nest-boxes pointing directly at the entrance, connected to Raspberry Pi 5 single-board computers (Raspberry Pi Ltd., Cambridge, UK). These modules feature a 12 MP Sony IMX708 image sensor and an infrared-sensitive lens, which performed well under the low-light conditions of the nest-boxes. In addition, these units supported high-speed video recording at 50 fps, allowing capture of events lasting as little as 1s. Cameras operated every one to three days from 7:00 to 8:30 each day, to capture peak morning provisioning activity. If any recordings failed, we manually restarted the cameras later in the day. We recorded broods ranging from six to 15 days old (mean = 11.7 ± 0.3 days) and filmed each brood on a median of 2 (± 1.6) days. In total, the recording effort resulted in 3,733 minutes of footage across 15 nest-boxes, yielding 38 videos of approximately 90 minutes each.

We manually analysed these videos in Boris software (version 9.2.3; Friard & Gamba, 2016) by continuously watching their full duration and annotating timestamps for nest-box entrances and exits. Entrances (“IN”) were recorded once a bird reached the inside of the nest chamber, exits (“OUT”) when a bird was seen leaving the chamber entrance.

### S8B. Great tit frame extraction from annotated videos

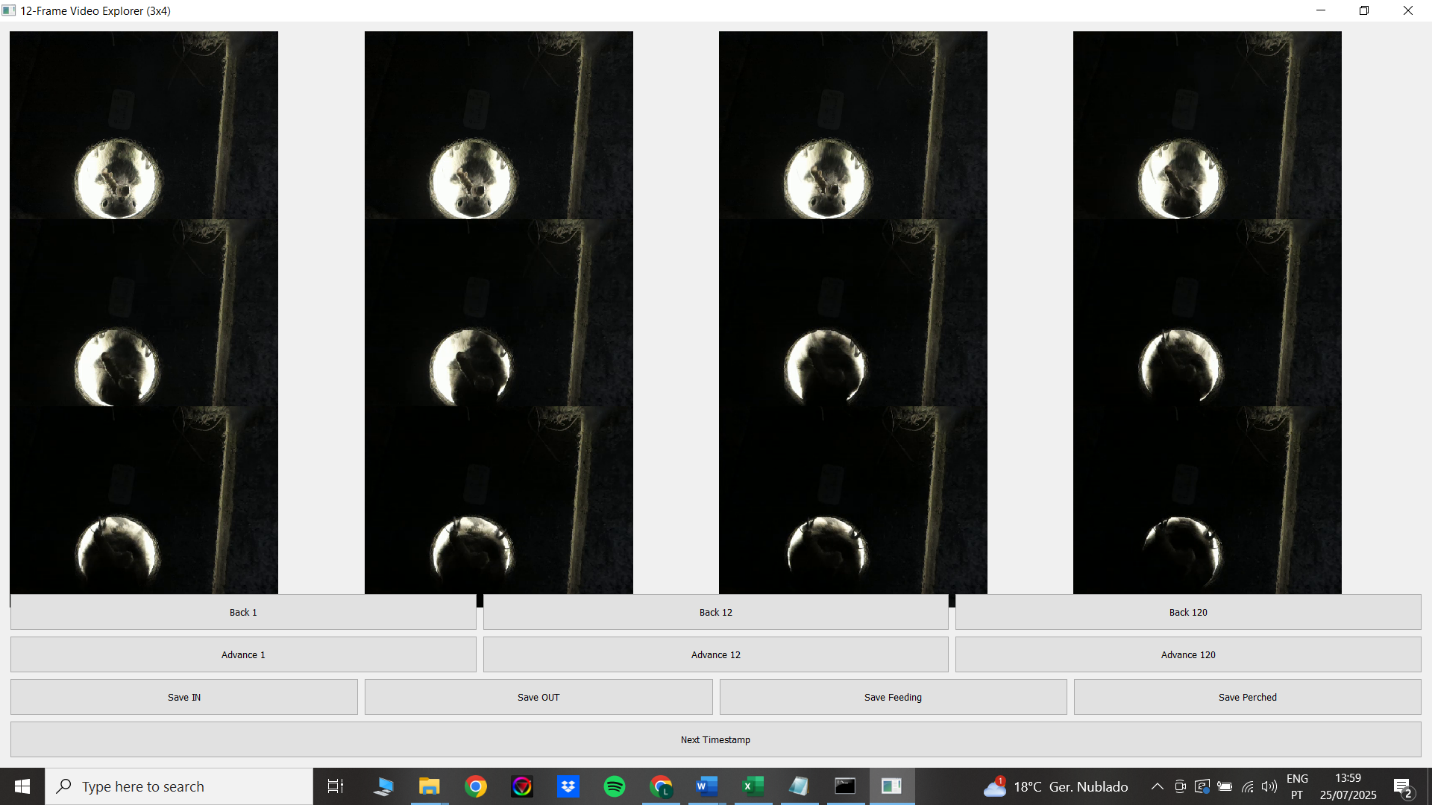
Similarly to the sociable weaver, great tit behavioural annotations were marked to the second and not by frame. Moreover, to find a specific sequenced frame interval to be used in the modelling stage, all annotations were screened by every 12 frames using a custom-built frame extraction interface, which allows saving already labelled sequences of frames for each defined behaviour (“IN”, “OUT”, Fig. S4). The same was not done for the sociable weavers, as the annotation dataset was too large for a unique screening of each event.

**Figure S7:** Custom-built frame viewer interface for fine-grained behavioural analysis and frame export. This Python application, developed with PyQt5 and OpenCV, allows loading a video and a corresponding file containing behavioural event timestamps. Users can view and scroll through 12 consecutive frames around each event and save selected frames into categorised folders (e.g., “IN”, “OUT”).

### S8C. Great tit LSTM modelling

We trained a LSTM model for great tits following the same procedures described in S5B. However, because data for great tits were very limited, we applied a leave-one-out approach, which means that we trained 10 different models instead of one. At each training iteration, one nest was excluded from training and used solely for validation. Accuracy and other performance metrics reports are therefore calculated from the combined predictions 10 different models, with each nest being predicted only by a model in which that nest was not included in the training dataset.

### S8D. Great tit LSTM evaluation

Our great tit behavioural automation was evaluated in the same way as for the other two species. However, the training dataset was relatively small. The models were tested on 10 videos from 10 different nests, comprising a total of 16.39 hours of footage and 789 recorded nest-box movements. Model processing speed was assessed directly on these videos. Model performance and speed were only assessed for the models, as for the blue tits. The only difference is that we did not apply any confidence threshold for the prediction. Based on preliminary analyses of 3 videos we concluded that even including a small confidence threshold as in the blue tits, would raise the production of false negatives without significantly decreasing the produced false positives.

### S8E. Great tit LSTM additional metrics

**Table S10:** For complete model evaluation, the Macro F1 score is reported for the great tit nest activity detection model. This metric represents the harmonic mean of precision and recall, computed per class and averaged equally across classes, with values closer to 1 indicating better performance.

^a^Nest-chamber entrance, ^b^nest-chamber exit, ^c^negative class

| *Model* | Classes | Macro F1 score |
| --- | --- | --- |
| *Nest activity* | IN^a^ vs. OUT^b^ vs. NC^c^ | 0.93 |

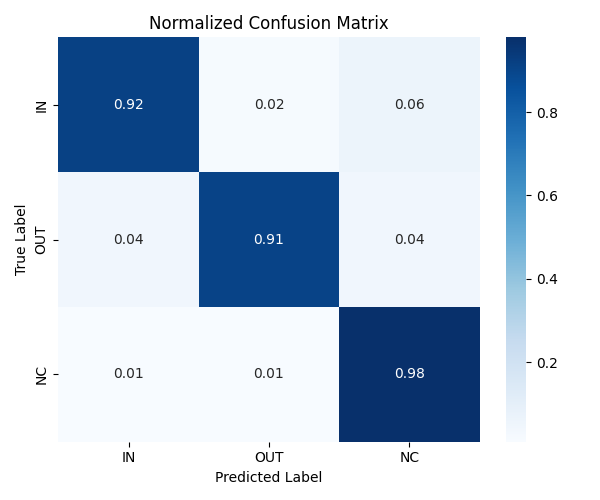

**Figure S8:** Normalized confusion matrix is presented for the great tit nest activity detection model, to illustrate classification performance. This matrix displays the proportion of predicted versus true classes, highlighting both correct classifications and misclassifications, and complement the Macro F1 evaluation by showing class-level performance.

### S9. Dataset properties and importance

Compared to datasets generated under controlled laboratory conditions, our dataset collected from wild bird species, presents a range of unique challenges and strengths that are worth highlighting. These aspects help to understand the context of our automation approach, but also to show the broader value and difficulty of behaviour classification in natural environments.

First, wild environments are inherently uncontrolled, even when using nest-box footage. Such recordings exhibit substantial variability in lighting and background complexity. For example, the appearance of observed vegetation may change over time, either from new nest material accumulating inside nest boxes or due to the naturally variable structure of sociable weaver nest chambers, where entrance shapes and visibility differ between chambers and even across days. Lighting conditions can fluctuate significantly throughout the day, affecting brightness and contrast. Additionally, dynamic elements such as wind can shift the position of the recording camera, while other birds may pass by or engage in unrelated behaviours within the frame. Other species are also common in this environment, such as predators, or insects either visiting the nest or appearing during feeding interactions when released by the parents.

Secondly, unlike behaviours observed in laboratory settings, which are often repetitive, constrained, and highly structured, our wild dataset captures animal behaviour that can be inherently irregular, context-dependent, and more difficult to standardise. This natural variability results in a wide range of behavioural expressions across individuals and situations, increasing the likelihood of ambiguity and hard behavioural classifications. In many cases, certain objects or actions may be visually confounded with behaviours of interest. For example, a bird flying over a sociable weaver nest chamber may be mistaken for one entering it; debris expelled from a chamber might resemble a bird exiting; or two birds arriving at or leaving a chamber simultaneously could be misinterpreted as an aggressive encounter. Similarly, in nest-box footage, a bird perching near the entrance but never entering may incorrectly be classified as participating in a feeding event but remains only looking. In the case of these specific ambiguous behaviours, our dataset, especially for the negative class (i.e., everything that is not the behaviour of interest), was hand-picked to include a big amount of such events, for a stronger model (see S6E). Additionally, recording in the wild occasionally captures rare and unpredictable behaviours (such as aggressive interactions among birds, infanticide, or predation events) that are biologically meaningful but difficult to document through manual annotation alone due to their rarity.

Finally, our dataset captures natural behaviours across extended timescales. For example, provisioning behaviour toward growing chicks inside a chamber can be observed over days or weeks. In the case of sociable weavers, this temporal depth is particularly valuable, as the dataset spans multiple breeding seasons and years. As a result, it incorporates a wide range of variation, seasonal and inter-annual differences, changes in weather conditions (including recordings from both sunny and rainy days), hardware differences (such as varying camera models and recording angles), and even individual-level variability, as the same birds may appear across different seasons. Additionally, differences among assistants in both recording setup and annotation style introduce further heterogeneity.

Together, these factors contribute to a uniquely rich dataset, with its diversity making it extremely valuable for further developing of other tools and models to study long-term behavioural patterns in nesting birds.

### S10. Detailed acknowledgments

From the Sociable Weaver Project, we thank Babette Fourie, Cecile Vansteenberghe, Franck Theron, Hugo Pereira, Marina Sentis, Marta Marmelo, Myriam El Harouchy, Sandra Esteves, Sophie Lardy, Rita Fortuna, Rita Leal and Zoe Tarren for their help in gathering the needed data that was the backbone of this work. Additionally, we thank De Beers Consolidated Mines for allowing us to work at Benfontein Reserve.

From the CEFE Tit Project, we thank Jérémie Moreau, Camille Bussière, and Hélène Dion-Phénix for their help with data collection in the field, Cristina Rios-Garcia for video analysis, and the teams of Centre d’Ecologie Fonctionnelle et Evolutive (CNRS) and of Evolutionary ecology of individual differences (UQAM) for support in organising the field work.

From the Wytham Tit Project, we thank Zoe Young and Grant Maslowski for their help during data collection, Ella Cole, Ben C. Sheldon and Keith McMahon for their advice in the conceptual and practical design of the work and their support in coordinating the field work.

Finally, we thank the Editors and Reviewers for their time and valuable feedback, which greatly improved this work.
